# FAK regulates IL-33 expression by controlling chromatin accessibility at c-Jun motifs

**DOI:** 10.1101/2020.07.15.177212

**Authors:** Billie G. C. Griffith, Rosanna Upstill-Goddard, Holly Brunton, Graeme R. Grimes, Andrew V. Biankin, Bryan Serrels, Adam Byron, Margaret C. Frame

**Affiliations:** Cancer Research UK Edinburgh Centre, Institute of Genetics and Molecular Medicine, University of Edinburgh, Edinburgh EH4 2XR, UK; Wolfson Wohl Cancer Research Centre, Institute of Cancer Sciences, College of Medical, Veterinary and Life Sciences, University of Glasgow, Glasgow G61 1QH, UK; Cancer Research UK Beatson Institute, Garscube Estate, Switchback Road, Glasgow G61 1BD, UK; MRC Human Genetics Unit, Institute of Genetics and Molecular Medicine, University of Edinburgh, Edinburgh EH4 2XU, UK

**Keywords:** FAK, c-Jun, IL-33, chromatin, transcription

## Abstract

Focal adhesion kinase (FAK) localizes to focal adhesions and is overexpressed in many cancers. FAK can also translocate to the nucleus, where it binds to, and regulates, several transcription factors, including MBD2, p53 and IL-33, to control gene expression by unknown mechanisms. We have used ATAC-seq to reveal that FAK controls chromatin accessibility at a subset of regulated genes. Integration of ATAC-seq and RNA-seq data showed that FAK-dependent chromatin accessibility is linked to differential gene expression, including of the FAK-regulated cytokine and transcriptional regulator interleukin-33 (*Il33*), which controls anti-tumor immunity. Analysis of the accessibility peaks on the *Il33* gene promoter/enhancer regions revealed sequences for several transcription factors, including ETS and AP-1 motifs, and we show that c-Jun, a component of AP-1, regulates *Il33* gene expression by binding to its enhancer in a FAK kinase-dependent manner. This work provides the first demonstration that FAK controls transcription via chromatin accessibility, identifying a novel mechanism by which nuclear FAK regulates biologically important gene expression.

## Introduction

Focal adhesion kinase (FAK) is a non-receptor tyrosine kinase that is overexpressed in many cancers, including squamous cell carcinoma (SCC) [1], breast, colorectal [2] and pancreatic cancer [3]. In addition to well-known localization at integrin-mediated cell-matrix adhesion sites (focal adhesions), FAK can localize to the nucleus, where it binds a number of transcription factors, including p53 [4], Gata4 [5] and Runx1 [6], to regulate the expression of *Cdkn1a* (which encodes p21), *Vcam1* and *Igfbp3*, respectively. By binding to these transcription factors, FAK has been linked to cancer-associated processes such as inflammation [5], proliferation [6] and survival [4].

Our previous work demonstrated that, in mutant H-Ras-driven murine SCC cells, nuclear FAK controls expression of cytokines and chemokines, for example *Ccl5*, to drive recruitment of regulatory T cells into the tumor microenvironment, resulting in suppression of the antitumor CD8^+^ T cell response and escape from anti-tumor immunity [7]. FAK regulates biologically important chemokines via its kinase activity and adaptor functions in the nucleus; briefly, nuclear FAK interacts with many transcription factors and accessory proteins in a gene expression-regulatory network [7]. Crucially, this includes the pro-inflammatory cytokine IL-33 that can either be found in the nucleus or be released from cells that are damaged or dying (alarmin) [8]. In SCC cells however, IL-33 is not secreted, but instead is translocated to the nucleus and functions there; in turn, nuclear IL-33 drives expression of immunosuppressive chemokines, such as *Ccl5* and *Cxcl10*, and we showed that IL-33 functions exclusively downstream of FAK in promoting pro-inflammatory gene expression and tumor growth [9]. However, the mechanisms by which FAK activity controls *Il33* gene expression are not understood, and this important extension of previous work is investigated here.

Our previous findings have suggested that FAK is present in the chromatin fraction and interacts with chromatin modifiers in the nucleus [7]. Therefore, we wanted to investigate whether FAK controls genome-wide chromatin accessibility changes and if these chromatin changes contribute to FAK-dependent gene expression, which to our knowledge has never been explored. To understand potential molecular mechanisms by which FAK regulates expression of genes like *Il33*, we examined FAK-dependent chromatin accessibility changes and transcription factor motif enrichment across the genome using ATAC-seq and integrated those data with RNA-seq. This revealed that FAK regulates chromatin accessibility of a subset of genes, and a number of these were differentially expressed in a FAK- and FAK kinase-dependent manner, including the previously identified FAK downstream effector *Il33*. Motif-enrichment analysis indicated that there was enrichment of sequence motifs known to bind ETS and c-Jun/AP-1 transcription factor family members on the *Il33* promoter and enhancer regions which are affected by FAK. Validation experiments confirmed that c-Jun is a key regulator of IL-33 expression in SCC cells by binding to the *Il33* enhancer, and that FAK’s kinase activity is important for regulating chromatin accessibility at this site. Analysis of genome-wide motif enrichment indicated that FAK likely regulates many more transcription factors beyond those already reported. Taken together, our data suggest that FAK is a common regulator of gene expression via modulating transcription factor binding to biologically relevant target gene promoters/enhancers by controlling chromatin accessibility, such as we demonstrate here using *Il33* as an exemplar. In turn, FAK-dependent gene expression changes, including *Il33*, are critically associated with cancer-associated phenotypes [7,9]. This is the first demonstration of how nuclear FAK can contribute to gene expression, and we report a new activity in the nucleus for a classical adhesion protein.

## Results

### FAK regulates transcription factor motif accessibility across the genome

FAK does not contain a DNA binding sequence and therefore likely regulates transcription indirectly, potentially through interactions with transcription factors and co-factors. To identify which transcription factors may be candidate mediators of FAK-dependent gene expression, we used ATAC-seq to analyze active (accessible) chromatin in FAK-deficient SCC cells or cells expressing different FAK mutant proteins. Specifically, we compared SCC cells in which FAK had been depleted by *Cre*-lox-mediated *Ptk2* (which encodes FAK) gene deletion (*FAK*^−/−^ cells) with the same cells re-expressing wild-type FAK (FAK-WT-expressing cells) or FAK mutants that were impaired in nuclear localization (FAK-nls-expressing cells) or deficient in kinase activity (FAK-kd-expressing cells) (cell lines generated previously [7,10]). These permitted the identification of FAK-dependent, FAK nuclear localization-dependent and FAK kinase-dependent alterations in chromatin accessibility and transcription factor motif enrichment in accessible regions of chromatin across the genome.

Chromatin accessibility analysis of ATAC-seq data detected 20,000–60,000 peaks (accessible regions) per sample (Supplementary Fig. S1A; see Supplementary Table S1 for further details of ATAC-seq statistics). The standard peak number in ATAC-seq experiments can vary depending on cell type, species, context and variations in the ATAC-seq protocol. Importantly, the peak number reported in our study is in the medium-to-high range for an ATAC-seq experiment performed in mouse cells (see additional file 2 in [11]).

The majority of peaks in FAK-WT-, FAK-nls-, FAK-kd-expressing as well as *FAK*^−/−^ cell lines were located 0–100 kb from the transcriptional start sites (TSS) (Supplementary Fig. S1B). The distance of ATAC-seq peaks from the TSS suggests that the accessible regions were predominantly located in likely enhancer [12] and promoter [13] regions.

We identified differentially accessible gene regions using the DiffBind package [14]. Differential peak calling was performed for each pairwise comparison for which FAK-WT samples were compared with each of the FAK knockout (*FAK*^−/−^) and FAK mutant (FAK-nls- and FAK-kd-expressing) cell lines. From this analysis, it was apparent that a subset of genes are regulated by FAK-dependent changes in chromatin accessibility (discussed further below).

We next analyzed the transcription factor motif sequences present in FAK-dependent differentially accessible peaks (hereafter termed motif-enrichment analysis). Motif-enrichment analysis allowed us to predict which transcription factors were regulating genes across the genome by analyzing the motif sequences in ATAC-seq peaks. We used the HOMER tool [15] to identify motif binding sites (i.e. genomic regions that match known transcription factor motifs) in the differentially accessible peaks identified by DiffBind analysis. Motifs in FAK-WT-enriched peaks were statistically compared to all the motifs identified in peaks called in the FAK-nls-, FAK-kd-expressing or *FAK*^−/−^ cells. This was expressed as a proportion of target sequences containing that motif (motifs in peaks enriched in FAK-WT-expressing cells) compared to the proportion of background sequences containing that motif (all motifs identified in the respective comparison, i.e. FAK-nls-, FAK-kd-expressing or *FAK*^−/−^ cells). This analysis detected multiple statistically significant changes in motif enrichment in differentially accessible peaks from FAK-WT vs FAK-deficient (*FAK*^−/−^) SCC cells (196 transcription factor motifs), FAK-WT- vs FAK-kd-expressing SCC cells (118 transcription factor motifs) and FAK-WT vs FAK-nls-expressing SCC cells (205 transcription factor motifs) (Benjamini–Hochberg-corrected *P* ≤ 0.05) (Supplementary Data S1). Importantly, the number of motifs identified in the ATAC-seq peaks were similar to those in previous published ATAC-seq datasets (see supplementary file 9 in [16]). These findings suggest that FAK regulates transcription factor motif enrichment in accessible regions of chromatin across the SCC genome.

In the motif-enrichment analyses of FAK-WT-expressing cells vs *FAK*^−/−^ cells and FAK-WT- vs FAK-nls-expressing cells, the two most highly enriched transcription factor motifs were for Jun–AP-1 and Fosl2 (all Benjamini–Hochberg-corrected *P* = 0), which exhibited an almost two-fold enrichment in motifs in the target (% target, FAK-WT-expressing cells) compared to the motifs identified in the background (% background, *FAK*^−/−^ cells or FAK-nls-expressing cells) (Fig. 1A). The top two hits in the motif-enrichment analysis of FAK-WT- vs FAK-kd-expressing cells were motifs for Ets1 and Etv1 (all Benjamini–Hochberg-corrected *P* = 0), which likewise revealed a two-fold enrichment in these motifs in the target (% target, FAK-WT-expressing cells) compared to the background motifs (% background, FAK-kd-expressing cells) (Fig. 1A). These data imply that FAK and specific FAK functions (kinase activity and nuclear localization) robustly regulate enrichment of particular AP-1 and ETS motifs within accessible chromatin regions in SCC cells.

**Figure 1.**
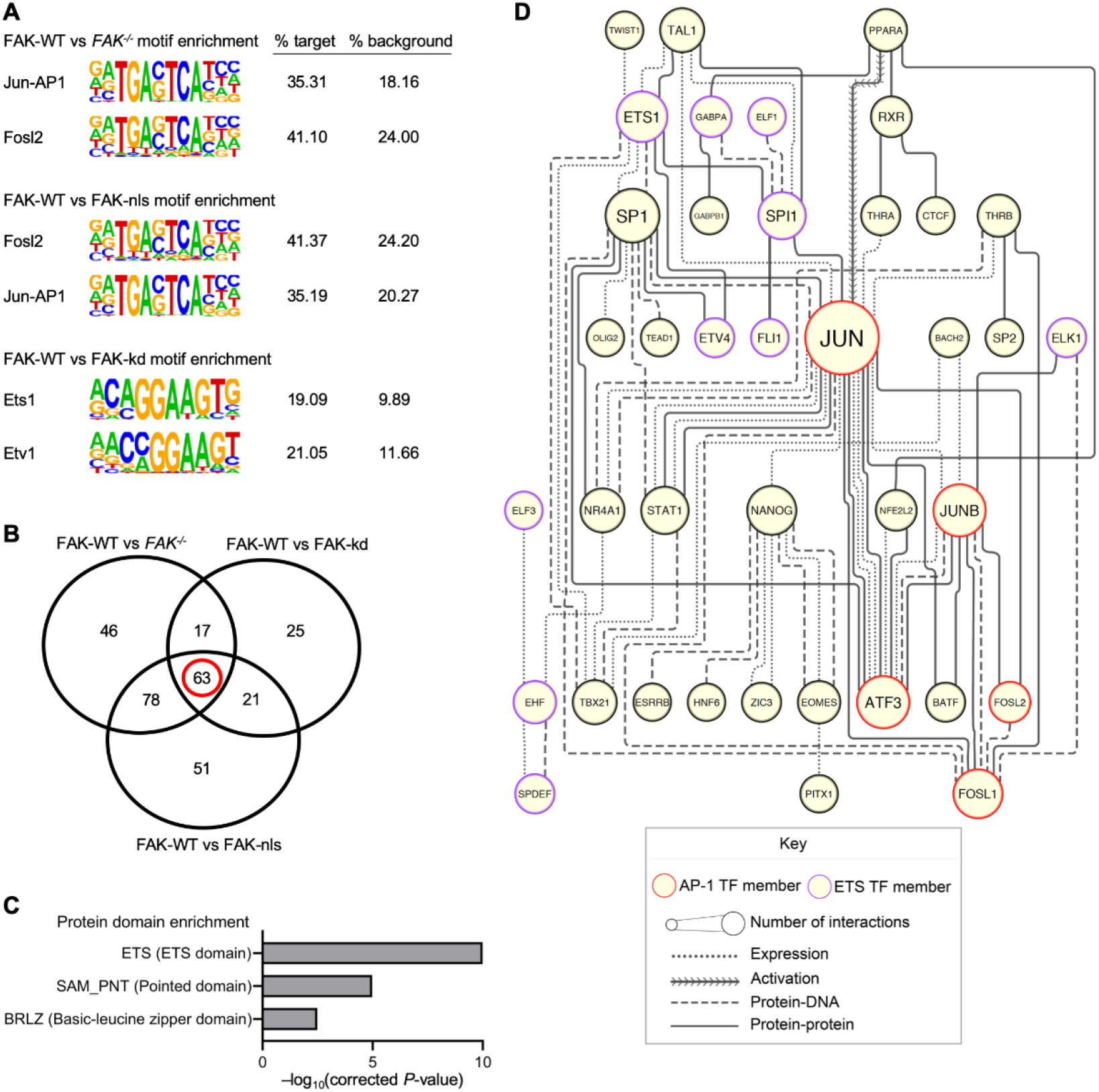
FAK regulates AP-1 and ETS motifs in accessible regions of chromatin. (**A**) The top two transcription factor motifs enriched in the FAK-WT-expressing vs *FAK*^−/−^ SCC cells (upper panel), FAK-WT- vs FAK-nls-expressing SCC cells (middle panel) and FAK-WT- vs FAK-kd-expressing SCC cells (lower panel) motif-enrichment analyses. Motifs in FAK-WT-enriched peaks (% target) were compared to all the motifs identified in peaks called in either the FAK-nls-expressing cells, FAK-kd-expressing cells or *FAK*^−/−^ cells (% background). The name of the transcription factor motifs are reported next to the consensus sequence (images from HOMER [15] published under a CC BY open access license), followed by the proportion of target sequences with that motif (% target) and the proportion of background sequences with that motif (% background). *P* ≤ 1 × 10^−31^ and *q* = 0 for all motifs shown. (**B**) Motif-enrichment data for FAK-WT-expressing cells vs *FAK*^−/−^ cells, FAK-nls-expressing cells or FAK-kd-expressing cells were filtered by set analysis to identify transcription factors enriched in FAK-WT-expressing cells but not enriched in *FAK*^−/−^, FAK-nls and FAK-kd motif-enrichment analyses (intersection set, red circle). (**C**) Protein domain-enrichment analysis of transcription factors associated with motifs that are enriched in FAK-WT-expressing cells. The full FAK-WT-expressing cells vs *FAK*^−/−^ cells motif-enrichment analysis (transcription factors predicted from both significant and non-significant motifs) was used as the background list. All terms with Benjamini–Hochberg-corrected *P* ≤ 0.05 are displayed (−log_10_-transformed corrected *P*-values are shown). The full domain name is reported in parentheses next to the corresponding SMART domain term. (**D**) Transcription factors that have enriched motifs in FAK-WT-expressing cells (intersection set in **B**, red circle) were used to construct a functional association network using Ingenuity Pathway Analysis. Only direct, mammalian interactions are shown. Edge (line) style represents type of physical or functional connection. Node (circle) size indicates the connectivity of the node (number of associations that node has within the network). Node borders for transcription factors from the AP-1 family are red and for the ETS transcription factors are purple. The network was structured using the yFiles hierarchical layout algorithm. *n* = 2 biological replicates.

We used set analysis to identify FAK-dependent transcription factors motif sequences in the ATAC-seq peaks (Supplementary Data S1). This analysis revealed enrichment of transcription factor motifs controlled by specific FAK functions (scaffolding, nuclear-localization and kinase activity). For example, in the FAK-WT vs *FAK*^−/−^ motif-enrichment analysis, there was enrichment of motifs known to primarily bind p53, which were not enriched in the FAK-WT vs FAK-nls or FAK-WT vs FAK-kd analyses (Supplementary Data S1), suggesting that FAK scaffolding functions may regulate exposure of p53 binding motifs. To establish the most relevant transcription factors responsible for FAK-WT-dependent gene expression, we filtered the transcription factors known to bind to FAK-regulated motifs that were only enriched in the FAK-WT-expressing cells when compared to *FAK*^−/−^ cells, FAK-nls-expressing cells and FAK-kd-expressing cells (63 transcription factor motifs; Fig. 1B and worksheet 4 in Supplementary Data S1).

To identify which transcription factors may regulate gene expression in the FAK-WT-expressing cells, we performed protein domain-enrichment analysis on the set of transcription factors known to bind FAK-regulated motifs (Fig. 1C). This analysis indicated that there was an over-representation of transcription factors known to bind motifs containing ETS (term SM00413:ETS, Benjamini–Hochberg-corrected *P* = 1.02 × 10^−10^) and PNT domains (term SM00251:SAM_PNT, Benjamini–Hochberg-corrected *P* = 1.06 × 10^−5^; Fig. 1C), including the ETS transcription factor family members Fli1, Elf3, Elf5, Gabpa, Spdef, Erg, Ehf and Ets1. Furthermore, there was also an enrichment for transcription factors known to bind motifs that contain basic-leucine zipper domains (term SM00338:BRLZ, Benjamini–Hochberg-corrected *P* = 0.0033; Fig. 1C), including members of the AP-1 family, such as c-Jun, JunB, Fosl1, Fosl2 and Atf3. Thus, our analyses revealed FAK-dependent enrichment of a set of sequence motifs known to bind AP-1 and ETS transcription factors.

To understand better the likely transcription factors responsible for FAK-dependent gene expression, we performed interactome analysis to determine putative connections between transcription factors known to associate with FAK-regulated motifs. We reasoned that transcription factors with exposed motifs in FAK-WT-expressing cells that have a large number of functional associations with other predicted transcription factors are more likely to be important mediators of FAK-dependent transcription. We constructed a functional association network, incorporating curated protein-protein and protein-DNA interactions, of the transcription factors whose motifs were enriched in FAK-WT-expressing cells (Fig. 1D). The network analysis revealed that transcription factors known to bind FAK-regulated motifs have a large number of connections with other transcription factors known to bind FAK-regulated motifs (Fig. 1D). The most highly connected transcription factor was the AP-1 member c-Jun (outlined in red in Fig. 1D), and network topology implied that c-Jun is a key signal integrator between all the other transcription factors in the network. Other well-connected nodes in the network were members of the AP-1 family, including JunB, Atf3 and Fosl1 (outlined in red in Fig. 1D). In addition, certain members of the ETS transcription factor family had many physical and functional connections within the network, namely Ets1 and Spi1 (outlined in purple in Fig. 1D). Collectively, these data suggest that FAK regulates motif enrichment in accessible regions of chromatin, in particular sequences known to bind to the AP-1 and ETS transcription factor family members.

### FAK regulates chromatin accessibility at a subset of genes, including Il33

Differential peak-calling analysis identified chromatin accessibility changes that were dependent on FAK, as well as FAK kinase activity and its nuclear localization (Fig. 2A). All ATAC-seq peaks were set to 500 bp to allow comparison between peaks in the SCC cell lines used in this study, and we reported distances from the ATAC-seq peak center as a heatmap (red indicates high read count (highly accessible region) in Fig. 2A). This analysis revealed ATAC-seq peaks across the genome with differential accessibility (varied read count) when comparing FAK-WT SCC cells to *FAK*^−/−^, FAK-nls-expressing and FAK-kd-expressing SCC cells, identifying changes in a subset of genes that varied depending on FAK status (Fig. 2A). These data implied that FAK regulates the chromatin accessibility at a subset of genes.

**Figure 2.**
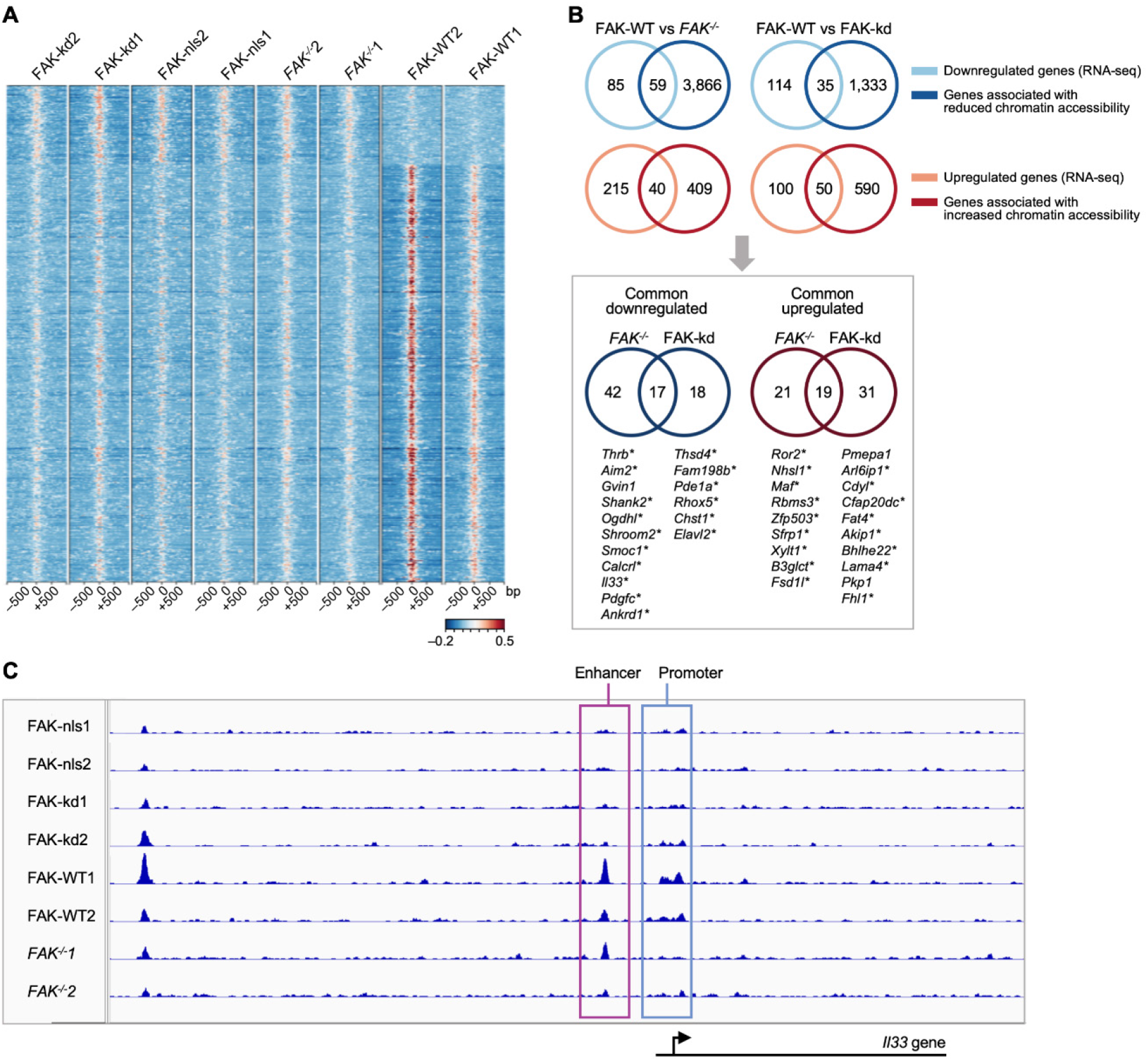
FAK regulates chromatin accessibility of a subset of genes, including the cytokine and transcriptional regulator *Il33*. (**A**) Heatmap representation of chromatin accessibility changes between FAK-WT, *FAK*^−/−^, FAK-nls and FAK-kd SCC cells (all peaks displaying differential accessibility are shown). Highly accessible regions (high read count, peak center) are shown in red, whereas less accessible regions (low read count) are indicated in blue. The *y*-axis of the heatmap reports the distance from the ATAC-seq peak center. Significantly different peaks were defined as those with a false discovery rate (FDR) ≤ 0.05. (**B**) The upper panel shows overlap of FAK-dependent (left) or FAK kinase-dependent (right) chromatin accessibility data with FAK-WT vs *FAK*^−/−^ or FAK-WT vs FAK-kd SCC RNA-seq differential expression data, respectively. Genes that were upregulated according to RNA-seq data were compared with genes exhibiting enhanced chromatin accessibility and vice versa. The lower panel (gray box) shows set analysis of common downregulated genes (left) and common upregulated genes (right) present in *FAK*^−/−^ and FAK-kd cells (with respect to FAK-WT cells), which also show changes in chromatin accessibility identified to be in common between FAK-WT vs *FAK*^−/−^ and FAK-WT vs FAK-kd analyses. In the lower panel, genes that display chromatin accessibility changes in FAK-nls cells in addition to *FAK*^−/−^ and FAK-kd cells are indicated by an asterisk. RNA-seq data for FAK-WT-expressing vs *FAK*^−/−^ SCC cells were reported previously [38] and re-analyzed here alongside the data for FAK-WT- vs FAK-kd-expressing cells. (**C**) Chromatin accessibility traces from FAK-WT, *FAK*^−/−^, FAK-nls and FAK-kd SCC ATAC-seq samples. Genomic regions that display FAK-dependent changes in chromatin accessibility upstream of the *Il33* gene are outlined in pink (enhancer) and blue (promoter region). Coverage is indicated on the *y*-axis, whereas genomic distance is shown on the *x*-axis. Below the chromatin accessibility traces is a schematic of the location of the *Il33* gene with respect to the ATAC-seq peaks. In (**A**) and (**C**), numbers appended to sample names indicate the biological replicate number for the respective cell line. *n* = 2 biological replicates for the ATAC-seq dataset (**A**–**C**); *n* = 3 biological replicates for the RNA-seq dataset (**B**).

We next identified which genes were regulated by FAK-dependent changes in chromatin accessibility using comparisons between the cell lines that varied only in FAK status. We wanted to determine which genes were associated with the ATAC-seq peaks enriched in FAK-WT-expressing cells (as identified by differential peak calling) to understand which genes are regulated by FAK-dependent accessibility changes. To assign each ATAC-seq peak to genes, we used ChIPseeker [17], which links each peak to the closest TSS using data from the University of California, Santa Cruz, genome browser annotation database (https://genome.ucsc.edu/). We used FAK RNA-seq data to confirm whether the genes that were regulated by FAK-dependent changes in chromatin accessibility were also differentially expressed in a FAK- and FAK kinase-dependent manner (Fig. 2B and FAK RNA-seq dataset reported in Supplementary Data S2). Set analysis identified genes that were either up- or down-regulated in a FAK- or FAK kinase-dependent manner, and also those genes whose FAK-dependent changes were associated with chromatin accessibility changes (intersection sets in upper panels in Fig. 2B). We found 36 genes whose expression and chromatin accessibility profiles were both regulated by FAK and its kinase activity (intersection sets in lower panel in Fig. 2B). Comparison of the FAK-nls mutant chromatin accessibility data to this subset of genes revealed that most of these are also dependent on FAK’s ability to localize to the nucleus (asterisks in lower panel in Fig. 2B).

As an exemplar, we next focused on one of these genes, *Il33*, because we had previously reported it as a FAK-regulated cytokine of biological significance in mediating FAK-dependent anti-tumor immunity [9]. Using ATAC-seq data to investigate whether chromatin accessibility was one mechanism by which FAK regulates *Il33*, we found that there were ATAC-seq peaks in FAK-WT-expressing SCC cells on *Il33* enhancer (−3199 from TSS) and promoter (+821 from TSS) regions (Fig. 2C). Moreover, these peaks were absent in the FAK-kd- and FAK-nls-expressing cells and reduced in *FAK*^−/−^ cells, which had no detectable peak on the promoter region and a suppressed ATAC-seq peak on the enhancer region. However, we note that the suppressed ATAC-seq peak on one replicate of the *FAK*^−/−^ cells (*FAK*^−/−^2) did not have sufficient read count to be identified as an ATAC-seq peak, and therefore the peak was not called. We conclude that FAK regulates chromatin accessibility at a subset of gene promoters, and some of these are differentially expressed in a FAK-dependent manner, as exemplified by *Il33*. This suggests that FAK-regulated, biologically important gene expression alterations may be controlled by FAK-dependent chromatin accessibility changes.

### FAK regulates IL-33 expression via chromatin accessibility at the c-Jun motif in the Il33 enhancer

In order to define key transcription factors that drive FAK-dependent *Il33* expression in mouse SCC cells, we performed motif-enrichment analysis on the ATAC-seq peaks proximal to the *Il33* promoter and enhancer regions in FAK-WT-expressing cells using HOMER (using data depicted in Fig. 2C). Analysis of the raw peak-calling data revealed that there was a number of peaks upstream of the *Il33* gene in the FAK-WT-expressing cells as well as *FAK*^−/−^ cells, FAK-nls- and FAK-kd-expressing SCC cell lines between −7480 and −42315 bp upstream of the TSS. To create a refined list to identify the key transcription factors important for *Il33* expression in FAK-WT-expressing cells, we excluded all the transcription factor motifs that were present in the aforementioned peaks upstream of the *Il33* gene in *FAK*^−/−^ cells, FAK-nls- and FAK-kd-expressing cells from our list of putative *Il33* transcription factor motifs. This identified 24 FAK-dependent transcription factor motifs, including sequence motifs known to be bound by the AP-1 components c-Jun, Atf2 and Atf7 (Fig. 3A and Supplementary Data S3).

**Figure 3.**
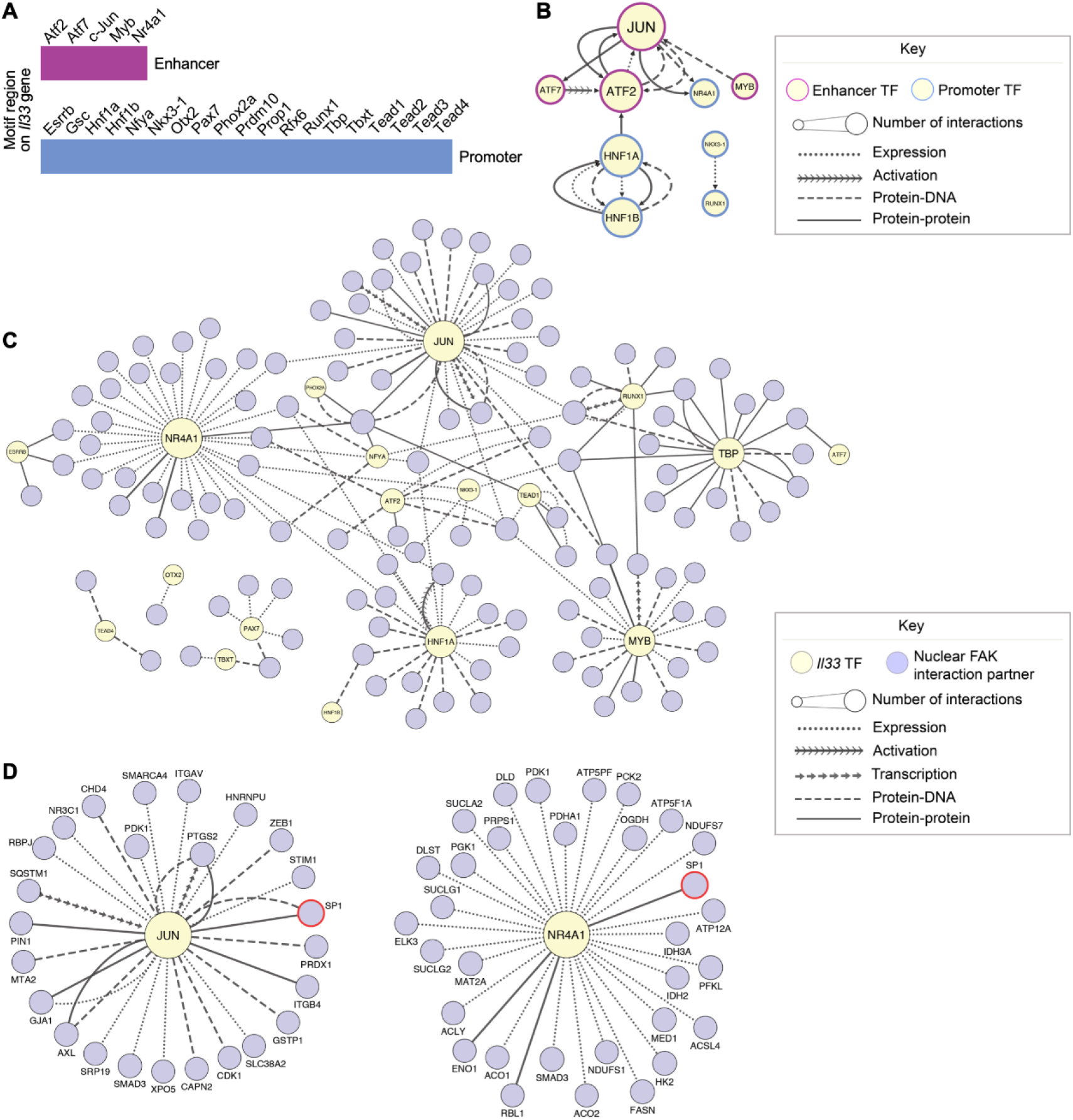
Interactome analysis identifies c-Jun as a key node in the *Il33* transcription factor network. (**A**) Set of transcription factors predicted to regulate *Il33* expression in FAK-WT SCC cells. Transcription factors that regulate *Il33* expression were predicted by analysing transcription factor motif sequences in the ATAC-seq promoter and enhancer peaks upstream of the *Il33* gene in FAK-WT-expressing cells using HOMER [15]. The potential *Il33* regulators and the respective location of the transcription factor motif upstream of the *Il33* gene (enhancer or promoter) are listed. (**B**) Functional association network of the predicted FAK-WT-enriched transcription factors on the *Il33* gene enhancer/promoter regions. Direct, mammalian connections are shown between the predicted transcription factors in FAK-WT-expressing cells. Node borders of predicted enhancer-associated transcription factors are pink; those predicted to bind the *Il33* promoter are blue. Node size indicates the connectivity of the node (number of functional connections that protein has within the network). The yFiles tree layout algorithm was applied to the network. (**C** and **D**) A previously published nuclear FAK interactome dataset [7] was integrated with the predicted transcription factors on the *Il33* promoter/enhancer regions. Upstream, mammalian interactions between the FAK nuclear interactors and the *Il33* transcription factors in FAK-WT-expressing cells were used to construct the network (**C**). The two most connected nodes, centered around c-Jun (left panel) and Nr4a1 (right panel) are detailed in **D**. Nodes for transcription factors predicted to bind to the *Il33* promoter/enhancer regions in FAK-WT-expressing SCC cell lines are colored in yellow; all potential FAK interactors that bind to predicted *Il33* motifs are colored purple (node labels omitted in **C** for clarity). Red node borders indicate proteins identified as FAK interactors by previous validation experiments [7,9]. Node size indicates the connectivity of the node. The yFiles organic layout algorithm was applied to the network. *n* = 2 biological replicates for the ATAC-seq dataset (**B**–**D**); *n* = 3 biological replicates for the proteomics dataset (**C** and **D**).

It is well established that in order for a transcriptional event to occur, transcription factors often need to form complexes with other transcription factors in the same or different families. For example, it is well known that c-Jun homodimerizes, as well as heterodimerizes with c-Fos and Fra or Atf family members, to regulate the expression of AP-1-dependent genes [18]. Furthermore, the transcription factor Nr4a1 has been shown to bind and co-operate with c-Jun to regulate the transcription of the *Star* gene [19]. Therefore, we addressed whether the transcription factors predicted to regulate *Il33* expression in FAK-WT-expressing cells can bind to and/or regulate each other. We reasoned that highly connected transcription factors may represent key nodes in the *Il33* transcription factor network and, therefore, potentially important regulators of IL-33 expression. Generation of an *Il33* transcription factor regulatory network for FAK-WT-expressing cells indicated that the transcription factors known to bind FAK-regulated motifs at the *Il33* gene have multiple functional connections (Fig. 3B). Indeed, the most highly connected transcription factor was the AP-1 member c-Jun (largest node in Fig. 3B), suggesting it may be a key node in the FAK-dependent *Il33* transcription factor network.

We next examined nuclear FAK binding partners (described previously in [7]) and used interactome analysis to contextualize these with regard to transcription factors that may bind to the identified sequence motifs in the *Il33* gene where accessibility is FAK-regulated. This enabled the prediction of putative *Il33* transcription factors that have functional connections with nuclear FAK binders and thereby may be regulated by FAK in a direct manner (Fig. 3C). The resulting network indicated that the transcription factors with motif sequences on the *Il33* promoter/enhancer have varying numbers of functional associations with putative nuclear FAK-interactors (indicated by node size in Fig. 3C). The transcription factors with the most links to nuclear FAK binding proteins were c-Jun and Nr4a1 (Fig. 3C). This implied that there were likely interesting connections between FAK and the transcription factors known to access motifs in the *Il33* promoter in a FAK-dependent manner. Our network analysis suggested that c-Jun interacts with a number of FAK binders identified previously [7], such as Pin1, which has been shown to bind to c-Jun and increase its transcriptional activity [20]. The FAK binding partner and transcription factor Sp-1 [9] has been reported to bind both the two most highly connected nodes in the network, c-Jun ([21]; left in Fig. 3D) and Nr4a1 ([22]; right in Fig. 3D). Other nodes that had connections with validated FAK binders included Tbp, which binds to the FAK binding protein Taf9 ([7]; Supplementary Fig. S2) to form the TFIID component of the basal transcription factor complex [23]. Therefore, our interactome analysis indicates that FAK is functionally well connected to transcription factors known to bind to sequence motifs on the *Il33* enhancer whose accessibility is FAK-regulated.

### c-Jun regulates IL-33 expression by binding to Il33 enhancer regions

Our nuclear FAK interactome analysis showed that c-Jun was a hub (i.e. a highly connected node) in the *Il33* transcription factor network (Fig. 3C). c-Jun is a component of the AP-1 family of transcription factors, and it is an important regulator of skin inflammation [24]. For example, c-Jun proteins are known to be important for the expression of the cytokine CCL5 [25], which we have shown to be regulated by FAK and IL-33, and loss of Jun proteins can lead to the onset of a chronic psoriasis-like disease [26]. Therefore, we hypothesized that c-Jun may be a likely regulator of inflammatory gene expression in SCC cells (which originate from skin keratinocytes) used in our studies. We performed siRNA-mediated depletion of *Jun* mRNA (which encodes c-Jun) (Fig. 4A), which led to a parallel significant downregulation of *Il33* mRNA (Fig. 4B) and reduced IL-33 protein expression (Fig. 4C). In addition, the FAK and IL-33 target gene in SCC cells, *Cxcl10*, also showed reduced mRNA levels as a result of *Jun* knockdown (Fig. 4D). Taken together, these data imply that c-Jun is likely an important regulator of IL-33 expression and of FAK- and IL-33-dependent target genes.

**Figure 4.**
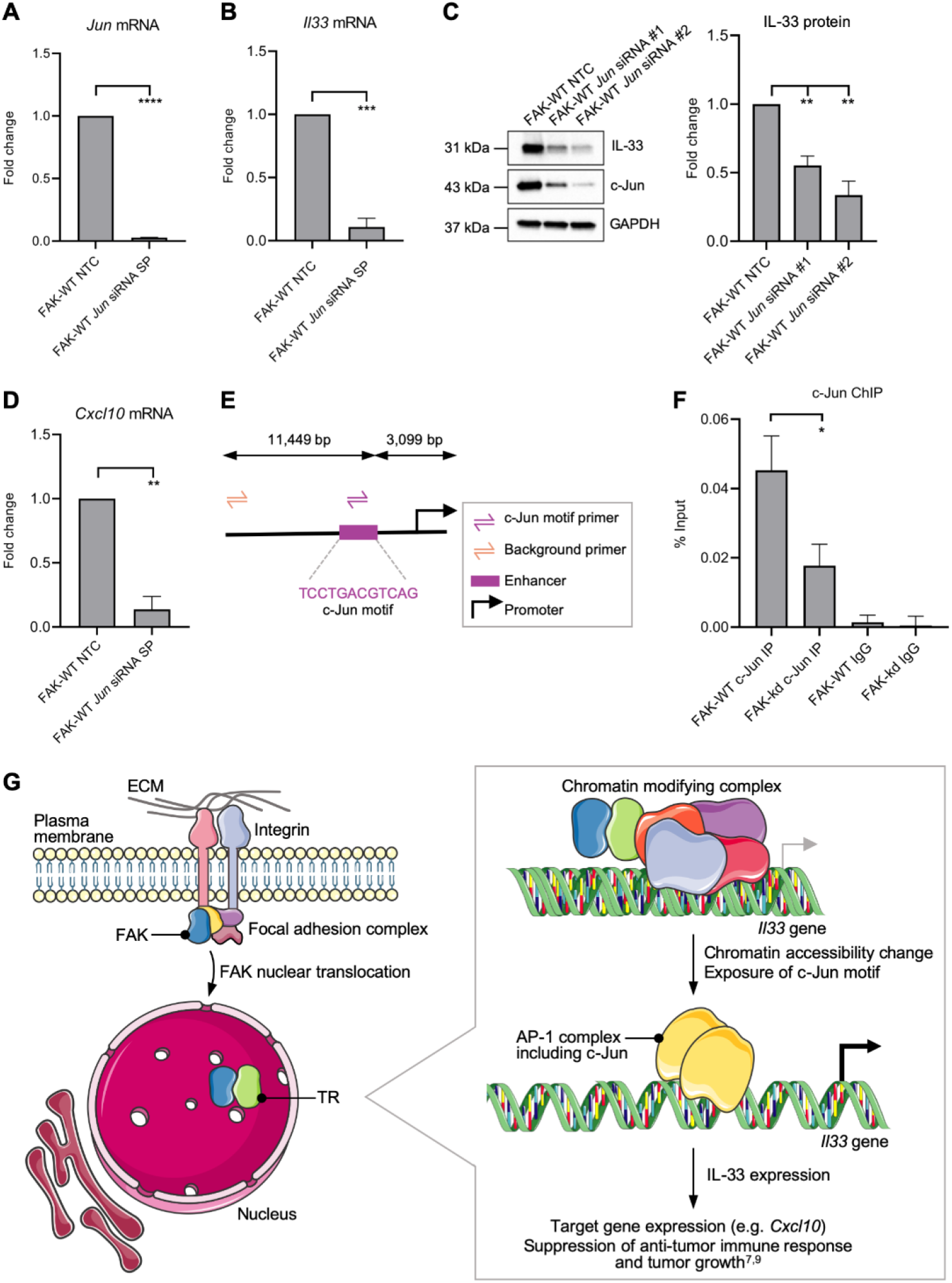
c-Jun regulates IL-33 expression by binding to *Il33* enhancer regions. (**A** and **B**) FAK-WT SCC cells were transfected with a non-targeting control (NTC) or *Jun* SMARTpool (SP) siRNA. *Jun* (**A**) and *Il33* (**B**) qRT-PCRs were carried out using *Jun* and *Il33* primers, respectively. Fold gene expression changes were calculated by normalizing cycle threshold (Ct) values to *GAPDH* and FAK-WT NTC Ct values. (**C**) FAK-WT-expressing cells were transfected with NTC or individual *Jun* siRNAs, and whole cell lysates were subjected to SDS-PAGE analysis. Blots were stained with IL-33, c-Jun and GAPDH antibodies (left panel). IL-33 protein expression was quantified by densitometry using ImageJ/Fiji software (v2.1.0, imagej.net/Fiji) and values normalized to GAPDH densitometry values (right panel). c-Jun knockdown was checked on a separate blot. Full length blots are reported in Supplementary Fig. S3. (**D**) FAK-WT cells were transfected with NTC or *Jun* SMARTpool siRNA. qRT-PCR was carried out using *Cxcl10* primers. (**E**) Schematic detailing the locations of ChIP primers upstream of the *Il33* gene. (**F**) Primers were designed to capture the c-Jun motif upstream of the *Il33* gene (c-Jun motif primer) and in the upstream region of the *Il33* gene to control for background binding (background primer). Pull-down efficiency was calculated using the % input method (see Materials and Methods). (**G**) FAK (blue) translocates to the nucleus and binds to transcriptional regulators (i.e. chromatin accessibility regulators and co-activators) (TR, green). At the level of the *Il33* gene, TRs potentially scaffold FAK to chromatin-modifying complexes to regulate chromatin accessibility changes at the *Il33* gene enhancer, allowing binding of the AP-1 complex containing c-Jun (yellow). AP-1 binding stimulates IL-33 expression, which suppresses the anti-tumor immune response and promotes tumor growth, as shown previously [9]. Images from Servier Medical Art (http://smart.servier.com/) were adapted under terms of a Creative Commons Attribution 3.0 Unported License: CC BY 3.0 Servier. Data are mean ± SEM. *n* = 3 biological replicates (**A**–**D**) or 5 biological replicates (**F**). ns, not significant; \**P* ≤ 0.05, \*\**P* ≤ 0.01, \*\*\**P* ≤ 0.001, \*\*\*\**P* ≤ 0.0001 by unpaired two-tailed *t*-test (**A**, **B** and **D**), one-way ANOVA (**C**) or *t*-test with Welch’s correction (**F**).

Next, we performed chromatin immunoprecipitation (ChIP)-qPCR analysis to confirm that c-Jun binds to the predicted c-Jun sequence-binding motif at the *Il33* enhancer, and whether or not FAK-dependent chromatin accessibility changes on the *Il33* enhancer are linked to perturbed c-Jun binding. Our HOMER [15] analysis identified a cAMP response element (CRE) (5’-TGACGTCA-3’) within the *Il33* enhancer peak, which are known to bind c-Jun–Atf dimeric complexes [18]. We therefore used ChIP to show that c-Jun binds to the CRE motifs at the *Il33* enhancer in a FAK-dependent manner via accessibility. Primers were designed around the region containing the CRE sequence motif in the *Il33* enhancer and an unrelated region upstream of this site to control for background binding (depicted in Fig. 4E). We used an anti-c-Jun ChIP-grade antibody to pull down DNA in formaldehyde-crosslinked, sonicated chromatin preparations from FAK-WT- and FAK-kd-expressing cells, since loss of FAK’s kinase activity displayed the most striking loss of chromatin accessibility at the *Il33* enhancer (Fig. 2C). Following immunoprecipitation, the DNA was purified and a qPCR was performed, whereby the *Il33* enhancer region and an upstream background region were amplified. We used the % input method to normalize the ChIP-qPCR data for potential sources of variability, including the starting chromatin amount in the chromatin extract, immunoprecipitation efficiency and the amount of DNA recovered (see Materials and Methods). Using ChIP, we found that c-Jun bound to the *Il33* enhancer in the FAK-WT-expressing cells (Fig. 4F). Furthermore, there was a significant loss of c-Jun binding in FAK-kd-expressing cells in comparison to FAK-WT-expressing cells (Fig. 4F). These data are consistent with FAK kinase activity regulating chromatin accessibility at the enhancer region upstream of the *Il33* gene at the predicted c-Jun binding site.

Next, we wanted to address whether FAK kinase activity may regulate the levels of phosphorylation of c-Jun. Transcriptional activation of c-Jun is mediated by phosphorylation of Serine 73 by c-Jun N-terminal kinase (JNK) [27]. Treatment of FAK-WT SCC cells with the FAK kinase inhibitor VS4718 resulted in a significant loss S73-c-Jun phosphorylation (Supplementary Figure S4). Interestingly, there was also a significant reduction of total c-Jun protein levels. There was therefore a change in the amount of cellular pS73-c-Jun upon treatment with a FAK kinase activity inhibitor, and we conclude that FAK kinase activity contributes to the amount of transcriptionally active c-Jun in the SCC cells used here.

Thus, we conclude that FAK, which is classically thought to be primarily an integrin adhesion protein, can function in the nucleus to control chromatin accessibility at specific gene promoters/enhancers. In turn, this leads to FAK-dependent transcription of specific genes, an example of which is the cytokine *Il33*. FAK/IL-33 downstream effectors significantly influence tumor biology [9].

## Discussion

In this study, we have discovered an undescribed function of nuclear FAK as a key regulator of chromatin accessibility changes and transcription factor binding. Furthermore, we have confirmed that nuclear FAK regulates c-Jun binding at the *Il33* enhancer region via chromatin accessibility changes to control *Il33* expression. As IL-33 is an important regulator of cytokine expression and tumor growth [8], FAK-dependent c-Jun regulation of IL-33 expression would be predicted to influence cancer cell biology, such as that we described previously [9]. It is perhaps not surprising that FAK can regulate c-Jun, since cytoplasmic-localized FAK is known to transduce signals through pathways such as MAPK [28] and Wnt [29,30], which are known to control c-Jun expression and its transcriptional activity [18,31]; however, what is surprising is the more direct link we have uncovered here between nuclear FAK function and its regulation of c-Jun transcriptional activity at the *Il33* enhancer via chromatin accessibility. Consistent with the links between nuclear FAK and c-Jun activity being more common, nuclear FAK is reported to regulate the expression of *Jun* (which encodes c-Jun) in response to ‘stretch’ in cardiac myocytes by binding to, and enhancing, the transcriptional activity of MEF2 [32].

Focal adhesion proteins other than FAK have been detected in the nucleus, such as Lpp [33] and Hic-5 [34], which are believed to function as transcription factor co-regulators [33,35]. Furthermore, the focal adhesion protein paxillin can also translocate to the nucleus [36], where it contributes to proliferation [37], and we believe that there are other integrin-linked adhesion proteins capable of translocating to the nucleus and functioning at the nuclear membrane or inside the nucleus [38]. Relevant to the work we present here, a number of consensus adhesome components containing LIM (Lin11–Isl1–Mec3) domains have been directly linked to the regulation of chromatin accessibility and dynamics. For example, Hic-5 can inhibit the binding of the glucocorticoid receptor to the chromatin remodelers chromodomain-helicase DNA binding protein 9 (also known as ATP-dependent helicase CHD9) and brahma (also known as ATP-dependent helicase SMARCA2), resulting in a closed chromatin conformation and transcriptional repression of a subset of glucocorticoid receptor target genes [35,39]. Also, paxillin can regulate proliferation-associated gene expression by controlling promoter–enhancer looping via nuclear interactions with the cohesin and mediator complex [37]. Taken together, these reports suggest that focal adhesion proteins in the nucleus are capable of scaffolding chromatin remodeling complexes to regulate chromatin structure and gene expression changes.

An unanswered question is the mechanism by which FAK controls chromatin accessibility at regulated genes. In this regard, our previous nuclear interactome proteomics revealed that FAK can interact with proteins known to regulate chromatin accessibility [7]. These include the Smarcc2 and Actl6a components of the BRG1/BRM-associated factor (BAF) complex [7], which have been shown to be recruited to target gene enhancers by AP-1 to regulate chromatin accessibility [40]. IL-33 is required for the chromatin recruitment of the Wdr82 component of the chromatin-modifying protein serine/threonine phosphatase (PTW/PP1 phosphatase) complex [9,41]. IL-33 binds to the Brd4 (bromodomain-containing protein 4) transcriptional coactivator [9], which is known to recruit the BAF complex to target genes [42]. The nuclear FAK binding protein Sp-1 also interacts with the BAF complex to facilitate its recruitment to specific promoters [43]. Therefore, there is abundant evidence of connections between FAK or FAK binding proteins (i.e. FAK–Sp-1, FAK–IL-33) and FAK-regulated transcription factors (e.g. AP-1) to chromatin accessibility factors, such as the BAF complex and PTW/PP1 phosphatase complex. It is likely that FAK, and proteins to which it binds, scaffold chromatin remodeling proteins at target genes, such as *Il33* as we describe here, in order to determine the state of chromatin accessibility, the binding of transcription factors like AP-1 and transcription (Fig. 4G). Once IL-33 expression is activated, FAK and IL-33 bind and co-operate to regulate target gene expression (e.g. *Cxcl10*) in the nucleus by interacting with a network of chromatin modifiers and transcriptional regulators [9].

One limitation to this study is that our ATAC-seq analysis was only performed in one species. However, we note that the HOMER tool [15] uses motifs from a number of sources, including human and mouse. Indeed, many of the transcription factor motifs are highly conserved between related organisms, with DNA binding profiles for human and mouse transcription factors being almost identical. This high homology makes the information about transcription factors specifically interchangeable between organisms [15]. Furthermore, many transcription factors have high protein sequence homology between species. For example, mouse c-Jun is 96% homologous to its human counterpart [44]. In addition, the information used to predicted the c-Jun/CRE motif upstream of the *Il33* enhancer originated from a human ChIP-seq dataset (K562-cJun-ChIP-Seq(GSE31477)). Therefore, our mouse c-Jun ChIP analysis at this CRE motif has experimentally confirmed that there is homology in AP-1 binding motifs between mouse and human. As such, we believe that the data presented in this manuscript are also applicable to human cell lines.

In summary, we have discovered a completely new paradigm for how FAK may regulate transcription in the nucleus, i.e. as a critical regulator of chromatin accessibility changes at biologically important target genes, such as *Il33* we show here. Translocation of FAK to the nucleus, where it can bind to factors that control chromatin accessibility, can therefore communicate extracellular cues to the gene transcription machinery in the nucleus by this route.

## Materials and Methods

### FAK SCC cell line generation

Generation of the FAK SCC cell model has been described previously [10]. Briefly, K14*Cre*ER *FAK^flox/flox^* in FVB mice were subjected to the dimethylbenz[*a*]anthracene/12-*O*-tetradecanoylphorbol 13-acetate two-stage cutaneous chemical carcinogenesis protocol. FAK deletion was induced by culturing excised SCC cells in 4-hydroxytamoxifen. FAK-WT, FAK kinase-deficient (FAK-kd; K454R) or FAK nuclear localization sequence-mutated (FAK-nls; R177A, R178A, R190A, R191A, R216A, R218A) constructs were stably expressed in a *FAK*^−/−^ clone using standard retroviral induction protocols (for further details, see [7,10]). Cells expressing FAK-WT and FAK mutant constructs were selected using 0.25 mg/ml hygromycin.

### Tissue culture

*FAK*^−/−^ and FAK-WT-, FAK-nls- and FAK-kd-expressing SCC cell lines were cultured at 37°C, 5% CO_2_, in 1× Glasgow minimum essential medium (MEM) supplemented with 10% fetal bovine serum, 1% MEM non-essential amino acids, 1% sodium pyruvate, 1% L-glutamine and 1% MEM vitamins (all from Sigma-Aldrich).

### ATAC-seq

ATAC-seq samples were prepared similarly to described previously [45]. The specific ATAC-seq protocol used in this study has been previously reported [46]. ATAC-seq data were aligned to the *Mus musculus* reference genome mm10 using the bcbio ATAC-seq pipeline [47]. Accessible regions (i.e. ATAC-seq peaks) were called from the BAM files using the MACS2 algorithm [48] with the following parameters: -B -broad -q 0.05 -nomodel -shift −100 -extsize 200 -g 1.87e9. Differentially accessible regions between the FAK-WT-expressing cells and the *FAK*^−/−^, FAK-nls-expressing and FAK-kd-expressing cells were identified by differential peak calling using the R/Bioconductor package DiffBind [14], where significantly different peaks were defined as those with a false discovery rate (FDR) of below or equal to 0.05. Motif-enrichment analysis was performed using HOMER [15] following default parameters. ATAC-seq peaks were assigned to genes using ChIPseeker[17] with the following parameters: tssRegion = c(−500, 2000), annoDb = “org.Mm.eg.db”, TxDb = TxDb.Mmusculus.UCSC.mm10.knownGene.

Set analysis of ATAC-seq data in Fig. 1B and Fig. 2B was performed using VENNy software [49].

### siRNA transfection

FAK-WT SCC cells were transfected with either siGenome *Jun* SMARTpool (cat. no. D-001210-02-05; Dharmacon), siGenome *Jun* siRNA #1, siGenome *Jun* siRNA #2 (cat. no. MQ-043776-01-0002; Dharmacon) or non-targeting control siRNA #2 (cat. no. D-001210-02-05; Dharmacon) diluted in 500 μl Opti-MEM Reduced Serum Medium with GlutaMAX (Gibco) to a final concentration of 33 μM. Transfections were carried out using Lipofectamine RNAiMAX transfection reagent (Invitrogen) following manufacturer’s instructions. Cells were incubated in transfection mixes for 48 hours before harvesting for RNA extraction or whole cell lysate preparation.

### Chromatin immunoprecipitation (ChIP)-qPCR

The ChIP-qPCR experiments were performed as described previously [50,51]. FAK-WT- and FAK-kd-expressing cells (4 × 10^6^) were plated on 10-cm dishes (Corning) and then, the following day, were formaldehyde crosslinked and fractionated as described in [50]. Sonication was carried out using a BioRupter (Diagenode) at high power for 10 minutes (30-second on/off cycles) to yield DNA fragments of 1,000–200 bp in size. The samples were then centrifuged at 13,000 rpm for 15 minutes at 4°C. The supernatant was collected and 50 μl reserved for an input sample. For immunoprecipitations, a protein A and protein G Dynabead mixture (1:1) (both from Invitrogen) was added to sonicated lysate along with 0.48 μg of c-Jun antibody (cat. no. 9165; Cell Signaling Technology) or equal amount of rabbit IgG (cat. no. 2729; Cell Signaling Technology). Immunoprecipitations were incubated at 4°C overnight with rotation, prior to washing with 1× RIPA 150 mM NaCl (50 mM Tris-HCl [pH 8], 150 mM NaCl, 1 mM EDTA, 0.2% SDS, 0.2% sodium deoxycholate, 1% Triton X-100), 1× RIPA 500 mM NaCl (50 mM Tris-HCl [pH 8], 500 mM NaCl, 1 mM EDTA, 0.2% SDS, 0.2% sodium deoxycholate, 1% Triton X-100) and twice with cold Tris-EDTA (TE) buffer (10 mM Tris-HCl [pH 8], 1 mM EDTA [pH 8]), before resuspension in 200 μl of ChIP direct elution buffer (10 mM Tris-HCl [pH 8], 300 mM NaCl, 5 mM EDTA [pH 8], 0.5% SDS). Input samples were made up to 200 μl with 1× RIPA 150 mM NaCl and 0.4% SDS. Both the input and immunoprecipitation samples were incubated for 5 hours at 65°C. The following day, the input and immunoprecipitation eluates samples were treated with 2 μl of RNaseA/T1 (Ambion) for 30 minutes at 37°C and 5 μl of 10 mg/ml proteinase K (Ambion) for 1 hour at 55°C. The DNA was then purified using the QIAquick PCR purification kit (Qiagen) following manufacturer’s instructions. ChIP and input DNA were amplified using the following primers: c-Jun motif/*Il33* enhancer, F:ACCCTGGAGTGTTCTTTGCA and R:TGCCTTCTGAAGCTTACTCGA; negative control region, F:ATGTGTGCTGTGTGTATGCC and R:ACATTAAGGGCAGGAGACGT. ChIP-qPCR analysis was performed using SYBR Green master mix (Thermo Scientific) following manufacturer’s instructions. The following cycling conditions were used: 98°C for 10 seconds, 30× (98°C for 10 seconds, 60°C for 1 minute and 72°C for 4 minutes) and 72°C for 5 minutes. The% input method was used for c-Jun ChIP data normalization, whereby the cycle threshold (Ct) values of the ChIP samples are divided by the Ct values of the input sample (starting amount of chromatin used for the ChIP). The input sample Ct value was first adjusted using the following equation: adjusted input = Ct of input sample − log_2_(dilution factor). Then the% input was calculated for the CRE c-Jun ChIP and the background control region upstream of the *Il33* gene using the following calculation: 100 × (PCR amplification factor)^(adjusted^ ^input−PCR^ ^Ct^ ^value(c-Jun^ ^ChIP))^. Then the% input of the CRE motif was subtracted from the% input of the background control region to determine the amount of enrichment of c-Jun binding to the CRE motif over the background control region.

### RT-qPCR

RNA extraction was performed using an RNeasy Mini kit (Qiagen) following manufacturer’s instructions. cDNA synthesis was performed using the SuperScript First-strand Synthesis System (Invitrogen) following the manufacturer’s random hexamers protocol. qRT-PCR analysis was performed using SYBR Green master mix (Thermo Scientific) following manufacturer’s instructions. The following cycling conditions were used: 98°C for 10 seconds, 30× (98°C for 10 seconds, 60°C for 1 minute and 72°C for 4 minutes) and 72°C for 5 minutes. Primers used were as follows: *Il33*, F:GGATCCGATTTTCGAGACTTAAACAT and R:GCGGCCGCATGAGACCTAGAATGAAGT; *Cxcl10*, F:CCCACGTGTTGAGATCATTG and R:CACTGGGTAAAGGGGAGTGA; *GAPDH*, F:CTGCAGTACTGTGGGGAGGT and R:CAAAGGCGGAGTTACCAGAG. *Jun* was amplified using pre-designed primers from Qiagen (cat no. QT00296541).

### Cell lysis and immunoblotting

Cell lysis and immunoblotting was performed exactly as described previously [9]. Antibodies used in this study were as follows: IL-33 (cat. no. BAF3626; R&D Systems), c-Jun (cat. no. 9165; Cell Signaling Technology), Phospho-FAK (Tyr397) (cat. no. 3283; Cell Signaling Technology), Phospho-c-Jun (Ser73) (cat. no. 3270; Cell Signaling Technology), GAPDH (cat. no. 5174; Cell Signaling Technology).

### FAK kinase inhibitor treatment

FAK-WT SCC cells were treated with 250 nM VS4718 (Selleckchem) for 24 hours, prior to cell lysis and immunoblotting.

### RNA-seq

RNA was extracted from FAK-WT, *FAK*^−/−^ and FAK-kd SCC cells using an RNeasy kit (Qiagen) following manufacturer’s instructions. To verify sample purity, the samples were run on a 2100 Bioanalyzer using the Bioanalyzer RNA 6000 pico assay (both Agilent). Samples that achieved an RNA integrity number (RIN) of 8 or above were considered suitable purity for sequencing. Samples were prepared for sequencing using the TruSeq RNA Library Prep Kit v2 (low-sample protocol) (Illumina) and paired-end sequenced using a HiSeq 4000 platform (Illumina) at BGI.

Transcript abundance was determined using the pseudoalignment software kallisto [52] on the mouse transcriptome database acquired using the kallisto index, implementing default parameters. Quality control was performed on the kallisto output using MultiQC software (https://github.com/ewels/MultiQC). Transcript abundance was summarized to gene level and imported into the differential expression analysis R package DESeq2 [53] using the R package tximport [54]. Genes which had zero read counts were removed prior to differential expression analysis.

Differential expression analysis was performed using DESeq2 using default parameters, where FAK-WT vs *FAK*^−/−^ and FAK-WT vs FAK-kd SCC cell line gene read counts were compared. The Wald test was used for hypothesis testing in DESeq2, and all *P*-values were corrected for multiple testing using the Benjamini–Hochberg method. Transcripts that acquired a Benjamini-Hochberg corrected P-value of 0.05 or below and a log_2_-transformed fold change in expression of ≥1 or ≤−1 were considered significantly different between the cell lines. RNA-seq results from FAK-WT and *FAK*^−/−^ cells were previously analyzed [38]; RNA-seq data from all cell lines (Supplementary Data S2) were deposited in the NCBI Gene Expression Omnibus (GEO) [55] (GEO series accession number: GSE147670).

### Protein domain-enrichment analysis

Protein domain-enrichment analysis was performed for SMART domains using DAVID [56,57]. All terms that acquired a Benjamini-Hochberg corrected P-value of below 0.05 were considered statistically significant.

### Network analysis

Network analysis was performed using Ingenuity Pathway Analysis (QIAGEN Inc., https://www.qiagenbioinformatics.com/products/ingenuitypathway-analysis). The following parameters were used for network construction: database sources (Ingenuity expert information, protein-protein interaction database, BioGrid, IntAct), direct interaction, experimentally observed, protein-protein and functional interactions, mammalian interactions only. All networks were exported into Cytoscape [58], and the NetworkAnalyzer plugin [59] was used to visualize the most connected nodes in the networks before applying yFiles layout algorithms (yWorks).

### Statistical analysis

Statistical analysis was performed using Prism 8 (GraphPad Software). All *P*-values below 0.05 were considered statistically significant.

## Acknowledgements

We thank the Glasgow Precision Oncology Lab (Wolfson Wohl Cancer Research Centre, University of Glasgow) for running DNA sequencing for the ATAC-seq experiments; Jayne Culley for assistance with ChIP optimization and discussions; Catherine Naughton and Ryu-Suke Nozawa for advice on ChIP experiments and providing the ChIP protocol.

## Author contributions

B.G.C.G. designed (with B.S.) and performed the laboratory experiments, and carried out the network analysis with supervision from M.C.F., B.S. and A.B. R.U.-G. performed the bioinformatic analysis of the ATAC-seq data. H.B. trained B.G.C.G. in ATAC-seq sample preparation. G.R.G. performed the bioinformatic analysis of the RNA-seq data. B.G.C.G., M.C.F. and A.B. wrote the paper. M.C.F and A.V.B provided project funding. All authors commented on the manuscript and approved the final version.

## Funding

The work was funded by Cancer Research UK (grants C157/A15703 and C157/A24837 to M.C.F.).

## Conflict of interest

The authors declare that they have no competing interests.

## Data availability

The RNA-seq data have been deposited in the Gene Expression Omnibus and are accessible through GEO series accession identifier GSE147670.

## Notes

https://www.ncbi.nlm.nih.gov/geo/query/acc.cgi?acc=GSE147670

https://www.ncbi.nlm.nih.gov/geo/query/acc.cgi?acc=GSE161022

### Competing Interest Statement

The authors have declared no competing interest.

