## Supplementary data for "FAK regulates IL-33 expression by controlling chromatin accessibility at c-Jun motifs"

### Table of contents

**Supplementary Figure S1.** ATAC-seq analysis of FAK SCC cell lines.

**Supplementary Figure S2.** Connections between potential nuclear FAK binders and the *Il33* regulator Tbp.

**Supplementary Figure S3.** Full length immunoblots for Figure 4C.

**Supplementary Figure S4.** FAK regulates pS73-c-Jun and c-Jun levels.

**Supplementary Figure S5.** Full length immunoblot images for Supplementary Figure S4.

**Supplementary Table S1.** ATAC-seq statistics.

**A**

| Sample name | Total ATAC-seq peaks |
|---|---|
| FAK-WT1 | 53,728 |
| FAK-WT2 | 39,056 |
| *FAK$^{-/-}$1* | 43,153 |
| *FAK$^{-/-}$2* | 35,256 |
| FAK-nls1 | 24,402 |
| FAK-nls2 | 24,974 |
| FAK-kd1 | 32,961 |
| FAK-kd2 | 45,956 |

**B**

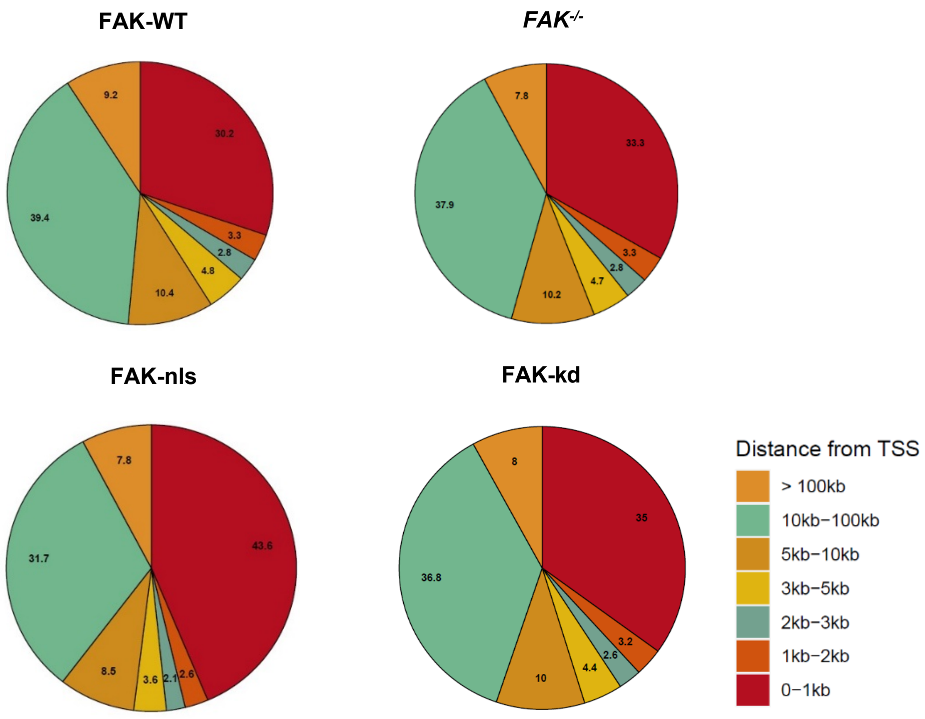


**Supplementary Figure S1. ATAC-seq analysis of FAK SCC cell lines. A.** Total number of ATAC-seq peaks identified for each replicate for each of the *FAK$^{-/-}$*, FAK-WT, FAK-nls and FAK-kd SCC cells. Numbers appended to sample names indicate the biological replicate number for the respective cell line. **B.** Pie charts indicating the mean relative distributions of ATAC-seq peaks for each cell line, binned by distance upstream from transcriptional start sites (TSS) of genes across the genome. *n* = 2 biological replicates.

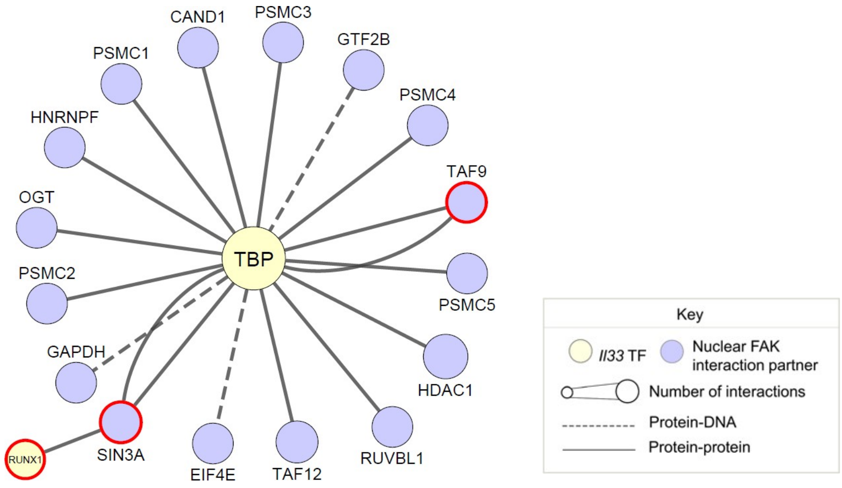


**Supplementary Figure S2. Connections between potential nuclear FAK binders and the *Il33* regulator Tbp.** A previously published nuclear FAK interactome dataset [7] was integrated with the predicted transcription factors on the *Il33* promoter/enhancer regions. Upstream, mammalian interactions between the FAK nuclear interactors and Tbp in FAK-WT-expressing cells were used to construct the network. The node for Tbp is colored yellow; nodes for all potential FAK interactors that bind to predicted *Il33* motifs are colored purple. Red node borders indicate proteins identified as FAK interactors by previous validation experiments [6,7]. The yFiles Organic layout algorithm was applied to the network. *n* = 2 biological replicates for the ATAC-seq dataset; *n* = 3 biological replicates for the proteomics dataset.

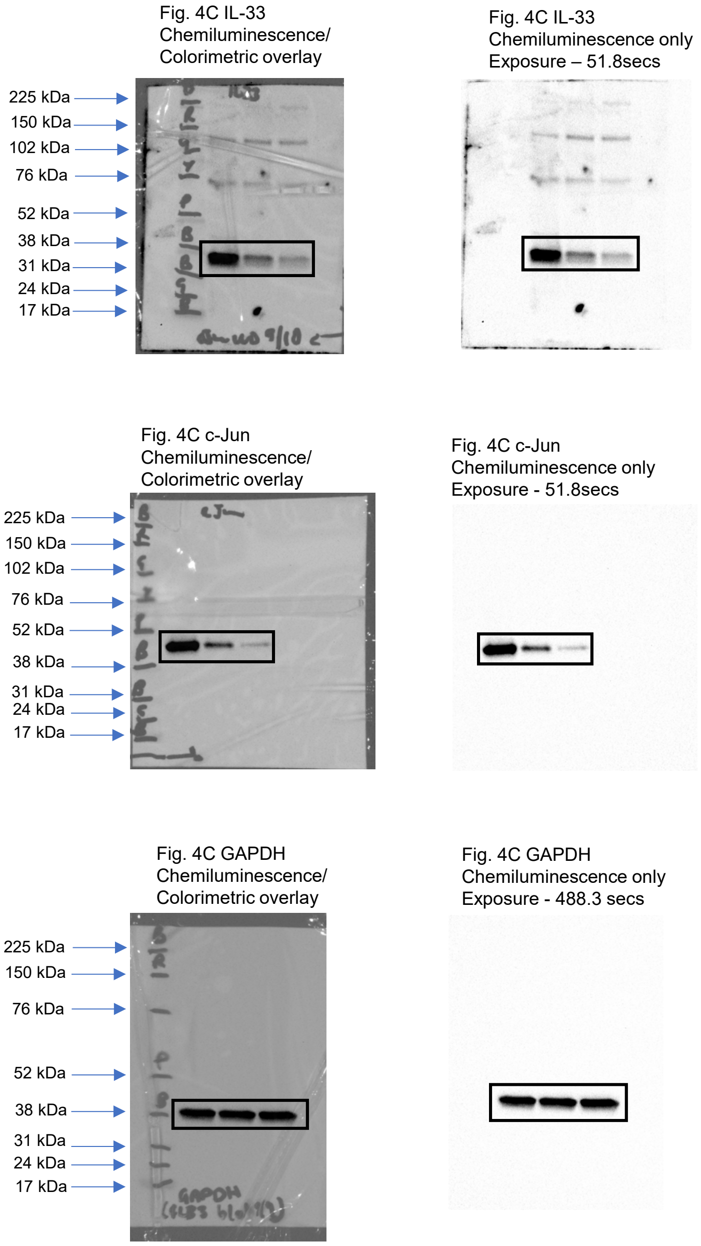


**Supplementary Figure S3. Full length immunoblot images for Figure 4C.** Images were taken using the Universal Hood 3 Bio-Rad Imager (Bio-Rad Laboratories). Chemiluminescence images were taken at various exposures (as indicated in text above right panels). Additionally, a colorimetric image was acquired to visualise protein markers, which was then overlaid onto the chemiluminescence image to confirm the protein bands on the gel match the molecular weight of the protein of interest. In left panel, chemiluminescence images are shown overlaid with the colorimetric images to show the location of the protein markers (highlighted with black marker written on blot) with respect to the protein bands (left panels). Chemiluminescence images of the full length blots are shown to the right panel. The specific panel in Figure 4C which the full length blots correspond to are indicated in text above the blots. Black rectangle highlights the protein band of interest. Amersham™ ECL™ Rainbow™ Marker - Full range (GE healthcare) was used for the protein ladder.

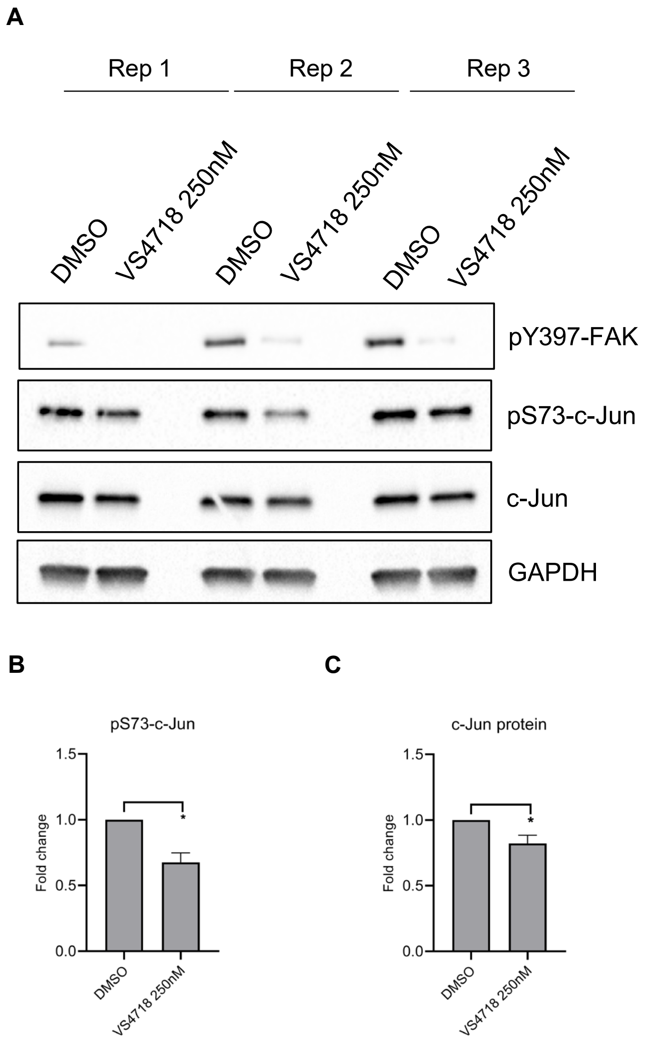


**Supplementary Figure S4. FAK regulates pS73-c-Jun and c-Jun levels. A.** FAK-WT-expressing cells were treated with 250nM VS4718 for 24 hours and whole cell lysates were subjected to SDS-PAGE analysis. Blots were stained with pY397-FAK, pS73-c-Jun, c-Jun and GAPDH antibodies. All three replicates shown (rep). The same samples were run on a separate blot to investigate effects of FAK kinase inhibition on total c-Jun expression. Full length blots are available in Supplementary Figure S5. **B.** pS73-Jun and **C.** Jun protein expression was quantified by densitometry using ImageJ/Fiji software (v2.1.0, imagej.net/Fiji) and values were normalized to GAPDH densitometry values. Unpaired t-test used to determine statistical significance. *n* = 3 biological replicates. * *P-value* ≤0.05

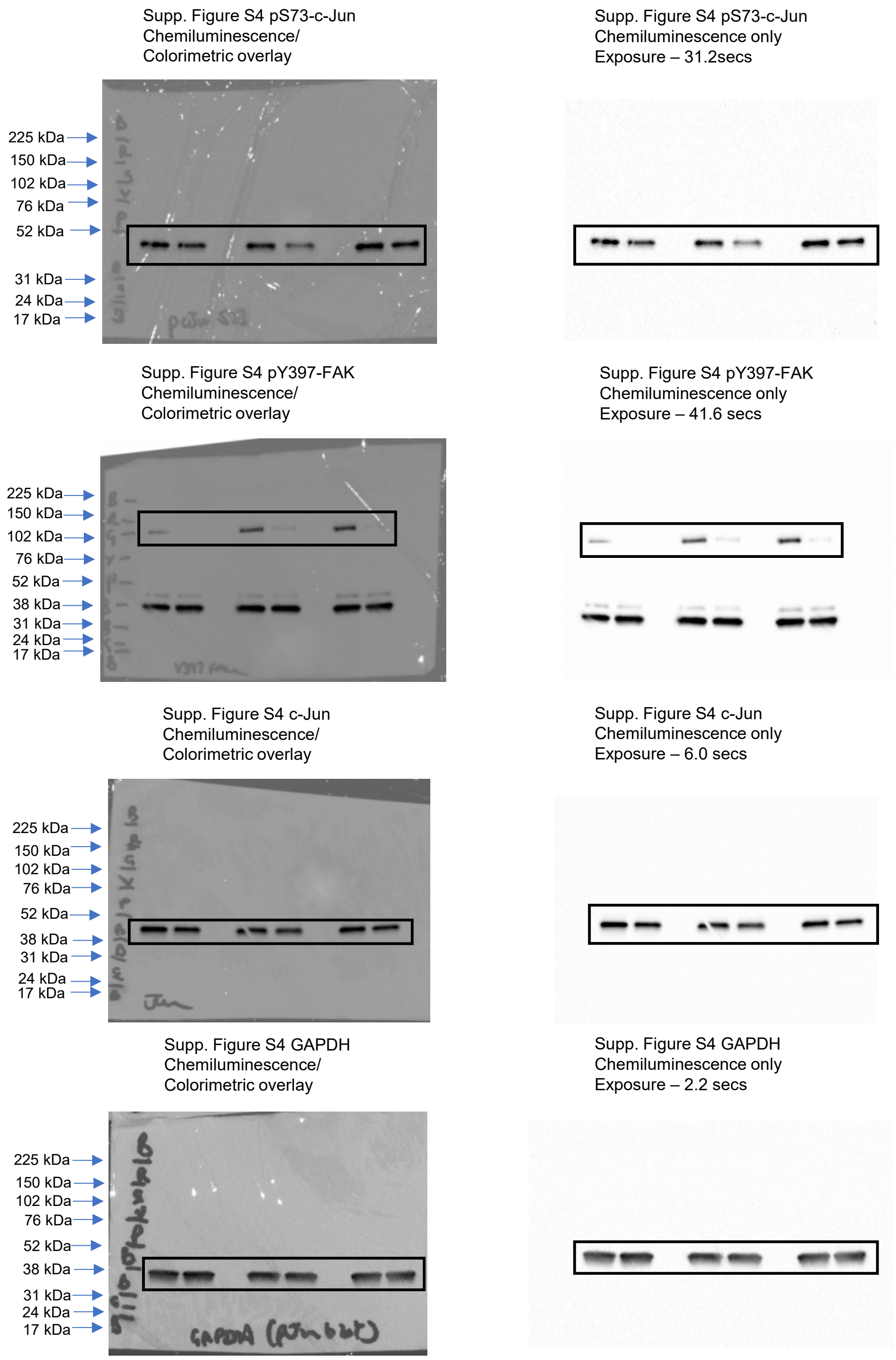


**Supplementary Figure S5. Full length immunoblot images for Supplementary Figure S4.** Images were taken using the Universal Hood 3 Bio-Rad Imager (Bio-Rad Laboratories). Chemiluminescence images were taken at various exposures (as indicated in text above right panels). Additionally, a colorimetric image was acquired to visualize protein markers, which was then overlaid onto the chemiluminescence image to confirm the protein bands on the gel match the molecular weight of the protein of interest. In left panel, chemiluminescence images are shown overlaid with the colorimetric images to show the location of the protein markers (highlighted with black marker written on blot) with respect to the protein bands (left panels). Chemiluminescence images of the full length blots are shown to the right panel. The specific panel in Supplementary Figure S4 which the full length blots correspond to are indicated in text above the blots. Black rectangle highlights the protein band of interest. Amersham™ ECL™ Rainbow™ Marker - Full range (GE healthcare) was used for the protein ladder.

| Sample | % duplications | Error rate | Reads mapped (×10$^6$) | Total seq. (×10$^6$) | % GC |
|---|---|---|---|---|---|
| FAK-WT1 | 32.80% | 1.92% | 82.9 | 130.9 | 46% |
| FAK-WT2 | 35.60% | 1.78% | 77.6 | 111.5 | 44% |
| *FAK$^{-/-}$1* | 31.80% | 1.94% | 82.2 | 131.0 | 44% |
| *FAK$^{-/-}$2* | 32.90% | 1.94% | 81.6 | 133.8 | 44% |
| FAK-nls1 | 36.70% | 2.05% | 65.0 | 112.9 | 46% |
| FAK-nls2 | 46.10% | 1.80% | 77.7 | 120.0 | 44% |
| FAK-kd1 | 20.20% | 2.14% | 57.7 | 101.5 | 46% |
| FAK-kd2 | 26.20% | 2.01% | 55.4 | 91.6 | 46% |

**Supplementary Table S1. ATAC-seq statistics.** General sequencing statistics of ATAC-seq analysis of FAK SCC cell lines. Total sequences (seq.) represent sequencing depth. % GC, proportion of guanine–cytosine content of the sequences. Numbers appended to sample names indicate the biological replicate number for the respective cell line.
